# Improvements to the Gulf Pipefish *Syngnathus scovelli* Genome

**DOI:** 10.1101/2023.01.23.525209

**Authors:** B Ramesh, CM Small, H Healey, B Johnson, E Barker, M Currey, S Bassham, M Myers, WA Cresko, AG Jones

**Affiliations:** Department of Biological Sciences, University of Idaho, Moscow, ID, 83844; Institute of Ecology and Evolution, University of Oregon, Eugene, OR, 97403; Presidential Initiative in Data Science, University of Oregon, Eugene, OR, 97403

**Author notes:** Contributed equally (WA Cresko, AG Jones).

## Abstract

The Gulf pipefish *Syngnathus scovelli* has emerged as an important species in the study of sexual selection, development, and physiology, among other topics. The fish family Syngnathidae, which includes pipefishes, seahorses, and seadragons, has become an increasingly attractive target for comparative research in ecological and evolutionary genomics. These endeavors depend on having a high-quality genome assembly and annotation. However, the first version of the *S. scovelli* genome assembly was generated by short-read sequencing and annotated using a small set of RNA-sequence data, resulting in limited contiguity and a relatively poor annotation. Here, we present an improved genome assembly and an enhanced annotation, resulting in a new official gene set for *S. scovelli*. By using PacBio long-read high-fidelity (Hi-Fi) sequences and a proximity ligation (Hi-C) library, we fill small gaps and join the contigs to obtain 22 chromosome-level scaffolds. Compared to the previously published genome, the gaps in our novel genome assembly are smaller, the N75 is much larger (13.3 Mb), and this new genome is around 95% BUSCO complete. The precision of the gene models in the NCBI’s eukaryotic annotation pipeline was enhanced by using a large body of RNA-Seq reads from different tissue types, leading to the discovery of 28,162 genes, of which 8,061 were non-coding genes. This new genome assembly and the annotation are tagged as a RefSeq genome by NCBI and thus provide substantially enhanced genomic resources for future research involving *S. scovelli*.

## Data Description

This article presents a resource (genome assembly) that marks a technological improvement compared to the one previously published in the article, “The genome of the Gulf pipefish enables understanding of evolutionary innovations.” [1].

A *de novo* genome assembly is evaluated based on three primary criteria: accuracy or correctness, completeness, and contiguity [2, 3]. Typically, the correctness of a genome is one of the most challenging features to measure. However, with modern, long-read sequencing technologies, the orientation of the contigs and the gene order of an assembly are highly accurate [4, 5, 6]. On the other hand, completeness and contiguity are easier to measure [6, 7, 8] yet more challenging to achieve, especially in non-model organisms. The Gulf pipefish (*Syngnathus scovelli*) genome is an essential resource for the study of comparative genomics, evolutionary developmental biology, and other related topics [1, 9, 10, 11, 12, 13, 14, 15]. Given the technological constraints when it was initially sequenced, the first version of the *S. scovelli* genome is highly accurate and mostly complete, but it leaves considerable room for improvement with respect to contiguity [1]. Here, with the use of third-generation sequencing technology, including PacBio High Fidelity (Hi-Fi) long reads from circular consensus sequences (CCS) and Hi-C proximity ligation from Phase Genomics, we produce a nearly complete chromosome-scale genome assembly that not only improves on completeness and accuracy but is also the most contiguous genome yet produced for the genus *Syngnathus* (Table 1).

**Table 1.**
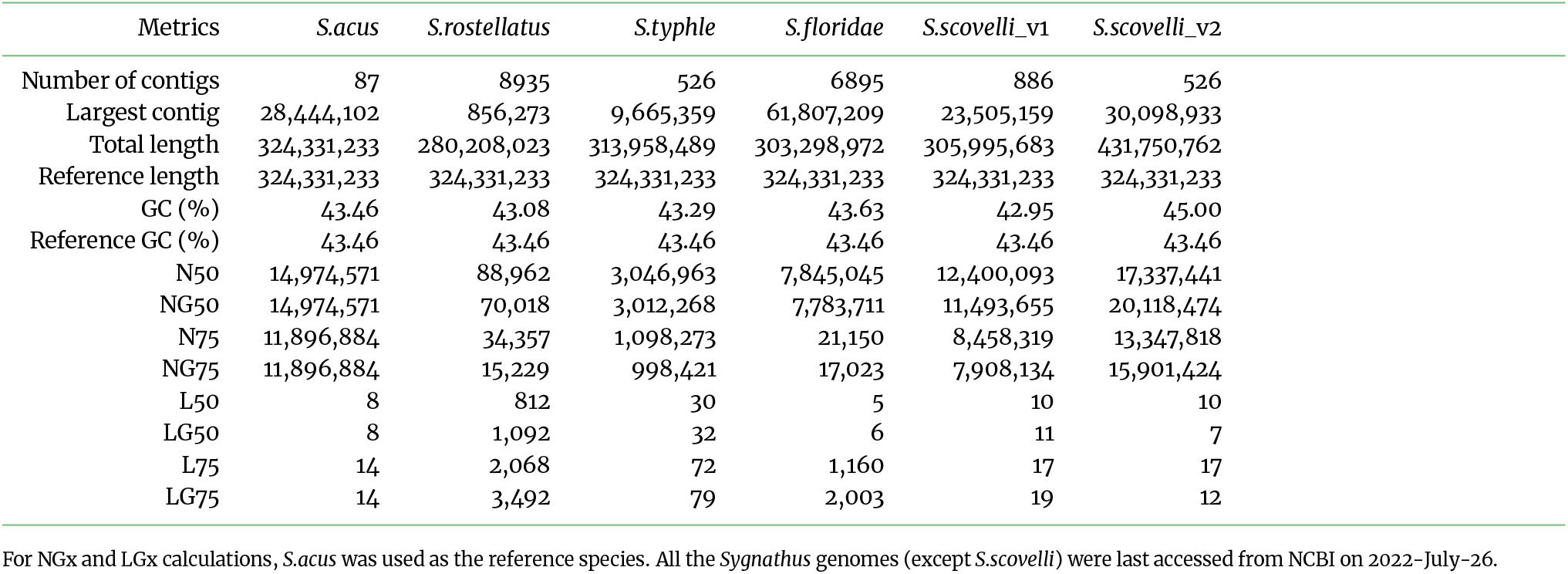
Contiguity metrics from QUAST for various *Syngnathus* species is summarized.

### Context

Evolutionary novelties are widespread across the tree of life. However, the origin of *de novo* genes and their associated regulatory networks, as well as their effects on the phenotype, remain mysterious in most species. Syngnathidae is a family of teleost fishes that includes pipefishes, seahorses, and seadragons [1, 12, 13, 14, 15, 17]. Syngnathid fishes are known for their evolutionary novelty with respect to morphology and physiology. For instance, species in this family have variously evolved elaborate leafy appendages, male brooding structures, prehensile tails, elongated facial bones, and numerous other unusual traits [1, 12, 13, 14]. With a variety of mating systems and sex roles [12, 13, 14, 15, 17], the syngnathid fishes also provide an excellent study system to investigate the generality of theories on sexual selection and reproductive biology [15, 17]. Advances in comparative genomics and the evolutionary developmental biology of novel traits in syngnathids will require the development of additional genomic tools. Among these are well-assembled and annotated genomes [1]. Here, we take a step in this direction by producing an improved reference genome for the Gulf pipefish.

## Methods

### DNA and RNA extraction

We collected *S. scovelli* from the Gulf of Mexico in Florida, USA (Tampa Bay), and flash froze them in liquid nitrogen. We pulverized approximately 50 mg of whole-body tissue (posterior to the urogenital opening) from a single male on liquid nitrogen, which we submitted to the University of Oregon Genomics and Cell Characterization Core Facility (UOGC3F) for high-molecular-weight DNA isolation using the PacBio Nanobind tissue kit. We submitted similar (but unpulverized) frozen tissue from the same individual fish to Phase Genomics to generate a Hi-C library using Proximo Animal (v4) technology.

In addition, we used organic extraction with TRIzol Reagent, followed by column-based binding and purification using the Qiagen RNeasy MinElute Cleanup Kit to extract mRNA from the Brain, Eye, Gills, Muscle/Skin, Testis, Ovary, Broodpouch, and Flap tissues.

### Sequencing and Assembly

After the size selection of genomic DNA using the Blue Pippin (11 kb cutoff), the UOGC3F constructed a sequencing library using the SMRTbell Express Template Prep Kit 2.0. One SMRT cell was sequenced by the UOGC3F using PacBio Sequel II technology, yielding 33.39 Gb in 2.05M CCS reads (out of 6.298M Hi-Fi reads in total). We sequenced 70.4 Gb of paired-end 150 nt reads (234.6 million in total) from the Hi-C library using an Illumina NovaSeq 6000 at the UOGC3F. The RNA sequencing libraries were prepared using KAPA mRNA HyperPrep Kit. We sequenced 159 bp paired-end reads using Illumina Novaseq 6000 for each tissue from the RNA sequencing libraries for annotation.

Using the Hi-Fi sequences, we estimated the genome size using genomescope2 (v2.0) [18] and meryl (v2.2) [19] with a default k-mer size of 21. The paired-end Hi-C reads were trimmed using trimmomatic (v0.39) [20] with the parameter HEADCROP:1 to remove the first base, which was of low quality. Together with the Hi-Fi sequences, we assembled the first-pass genome assembly in Hi-C integrated mode using hifiasm (v0.16.1) [19] with default parameters. The First-Pass assembly refers to the first draft consensus assembly from the Hi-Fi and Hi-C data. We extracted the consensus genome from hifiasm in fasta format and assembled the contigs into scaffolds using juicer (v1.6) [21]. We used the 3D-DNA (version date: Dec 7, 2016) [22] pipeline to merely order the scaffolds. The Hi-C contact map of the ordered scaffolds was visualized using juicebox (v1.9.8) with no breaking of the original contigs.

### Assessment of completeness and contiguity

To compare the completeness and contiguity of the latest version of the *S. scovelli* genome against the other *Syngnathus* genomes (Figure 1), we downloaded the genome assemblies of *S. acus* (GCA_024217435.2), *S. rostellatus* (GCA_901007895.1) [23], *S. typhle* (GCA_901007915.1) [23], and *S. floridae* (GCA_010014945.1) from NCBI. We used Benchmarking Universal Single-Copy Orthologs (BUSCO v5.2.2) [24] in genome mode with the actinopterygii_odb10 database (as of 2021-02-19) to evaluate the completeness of the genome. Also, we used a k-mer-based assessment using Merqury (v2020-01-29, [25]) to estimate the completeness and the base error rate. We then used the Quality Assessment Tool (QUAST v5.0.2) [26] to estimate Nx and Lx statistics for our assembly.

**Figure 1.**
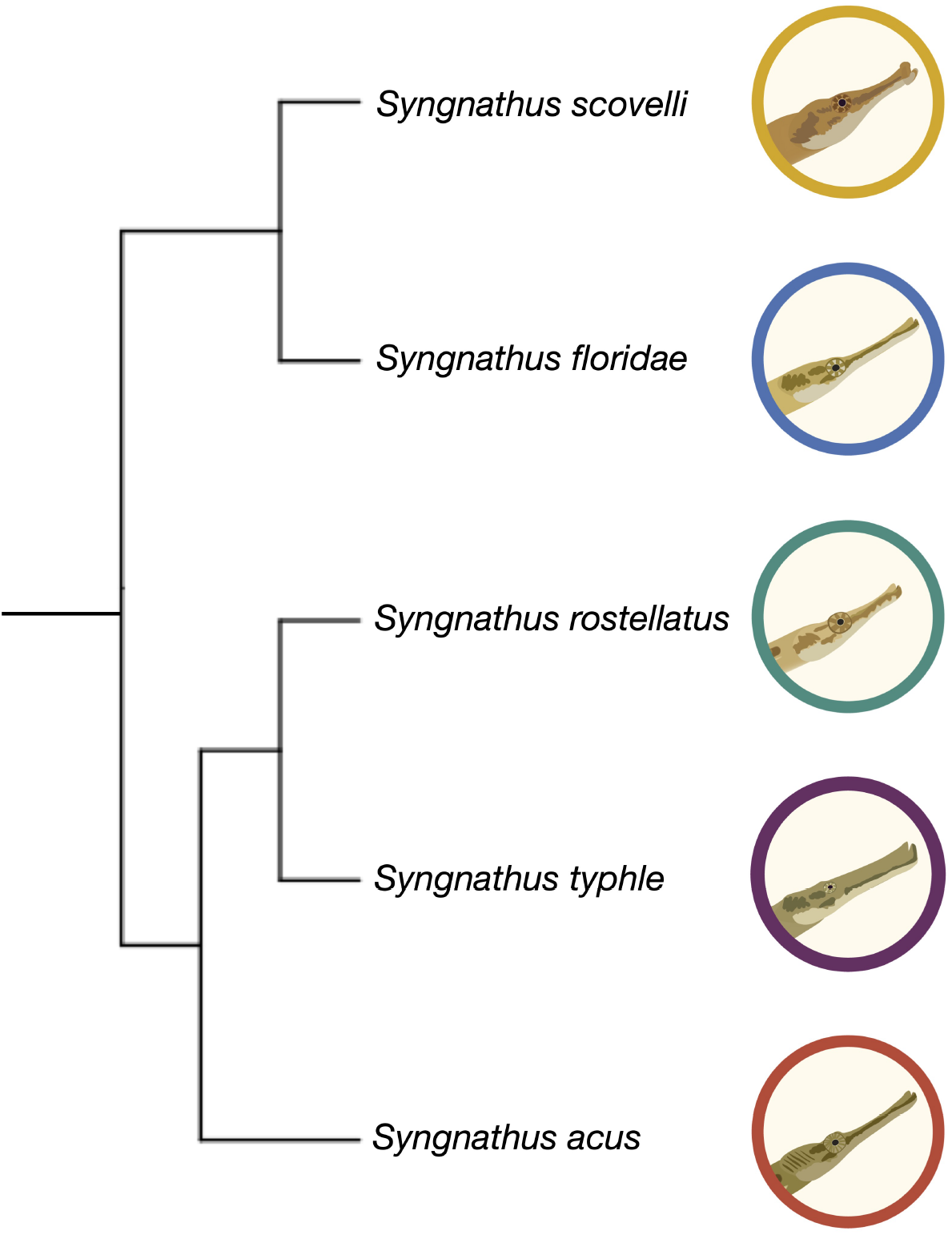
Cladogram of the five *Syngnathus* species in this study. This phylogeny is based on the Ultra Conserved Elements among all syngnathids [16].

### Annotation using the NCBI Eukaryotic Annotation Pipeline

The NCBI Eukaryotic Genome Annotation Pipeline (v10.0) is an automated software pipeline identifying coding and non-coding genes, transcripts, and proteins on complete and incomplete genome submissions to NCBI. The core components of this pipeline are the RNA alignment program (STAR and Splign) and Gnomon, a gene prediction program. In this pipeline, the RNA-Seq reads from the various (Brain, Eye, Gills, Muscle/Skin, Testis, Ovary, Broodpouch, and Flap) tissues of multiple samples, including the *S*.*scovelli* individual used for Hi-Fi and Hi-C sequence data (SRR20438584 - SRR20438604) were aligned to the genome. Gnomon combines the information from alignments of the transcripts and the *ab initio* models from an HMM-based algorithm to create a RefSeq annotation. This RefSeq annotation produces a non-redundant set of a predicted transcriptome and a proteome that can be used for various analyses. The Eukaryotic annotation pipeline is not publicly available; thus, we requested the staff at NCBI to annotate the *S*.*scovelli* genome.

## Data Validation and Quality Control

### Assembly Statistics

With approximately 2 million Hi-Fi reads and 234.6 million Hi-C reads, we generated the first pass consensus assembly with 585 contigs. The N50 and L50 for this assembly were 15.5Mb and 11, respectively. We scaffolded this assembly to correct misassembles and produced a final assembly containing 526 contigs with N50 and L50 values of 17.3 Mb and 10, respectively (Table 1). This improved version of the *S. scovelli* genome has around three times a lower number of contigs compared to the original *S. scovelli* genome. The NG50 and NG75 are ∼1.75x and ∼2x larger, respectively, than the previous assembly, implying less fragmentation. This new assembly reduces the number of gaps per 100 kilobase pairs (kb) from 6837.20 Ns per 100 kb to a mere 0.27 Ns per 100 kbp, owing to the increased contiguity. This new *S. scovelli* genome is on par with the current best genome in the *Syngnathus* genus, that of *S. acus*, which is a complete chromosome-scale assembly. The first 22 scaffolds of the *S. scovelli* genome are of chromosome-scale in line with the genetic map [1] and the karyotype data [27] with a total length of around 380Mb (Figure 3), comparable to the estimated genome size of 380 Mb (see supplementary details; Supplementary Table 1 and Figure 1). In addition, 88.94% of the total assembly length is captured in the 22 chromosome-scale scaffolds. For 15 of the chromosome-scale scaffolds, a single contig made up the total length; the remaining seven were generally composed of a small number of contigs (Figure 3).

**Figure 2.**
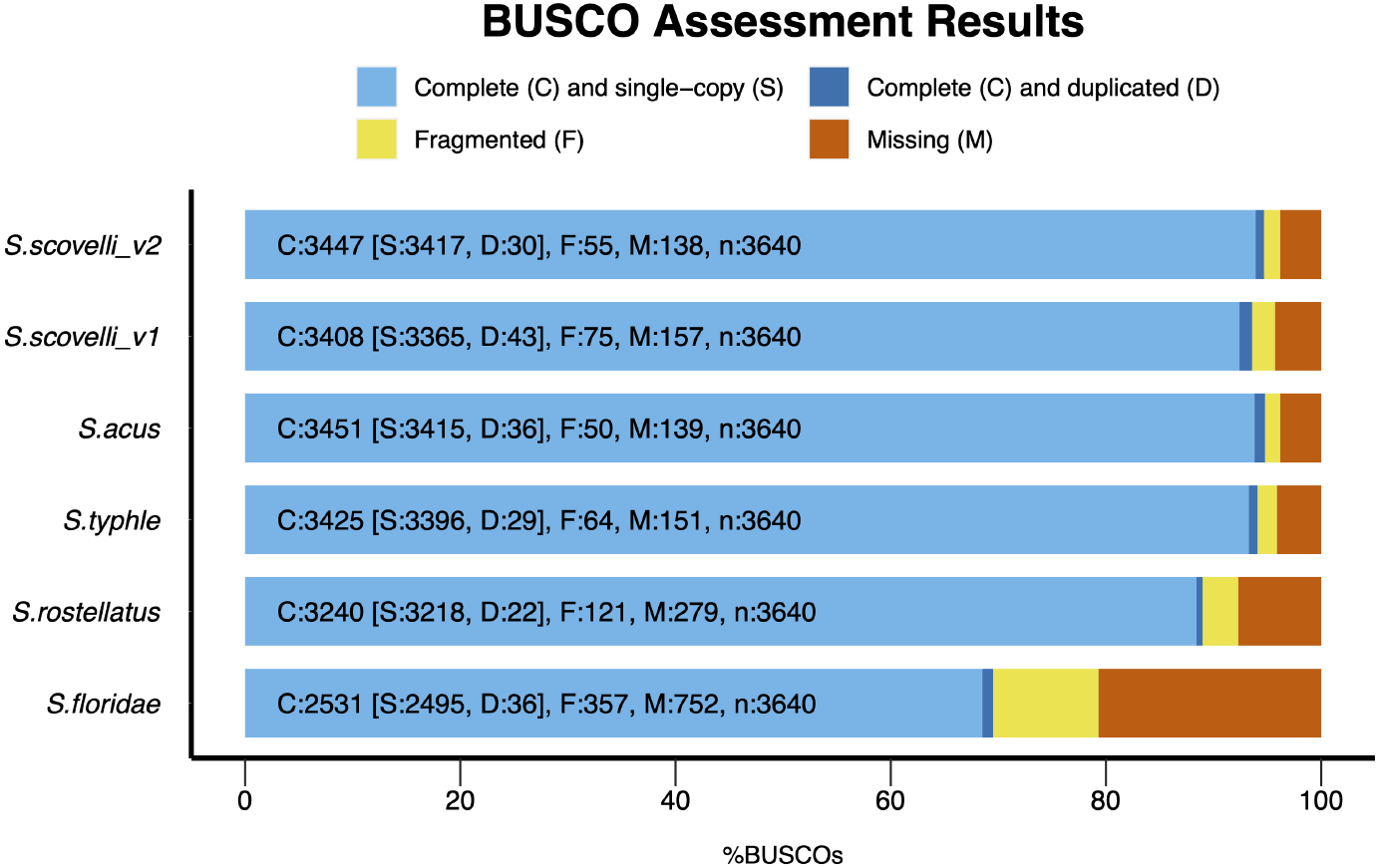
Comparison of BUSCO completeness among all the five *Syngnathus* species.

**Figure 3.**
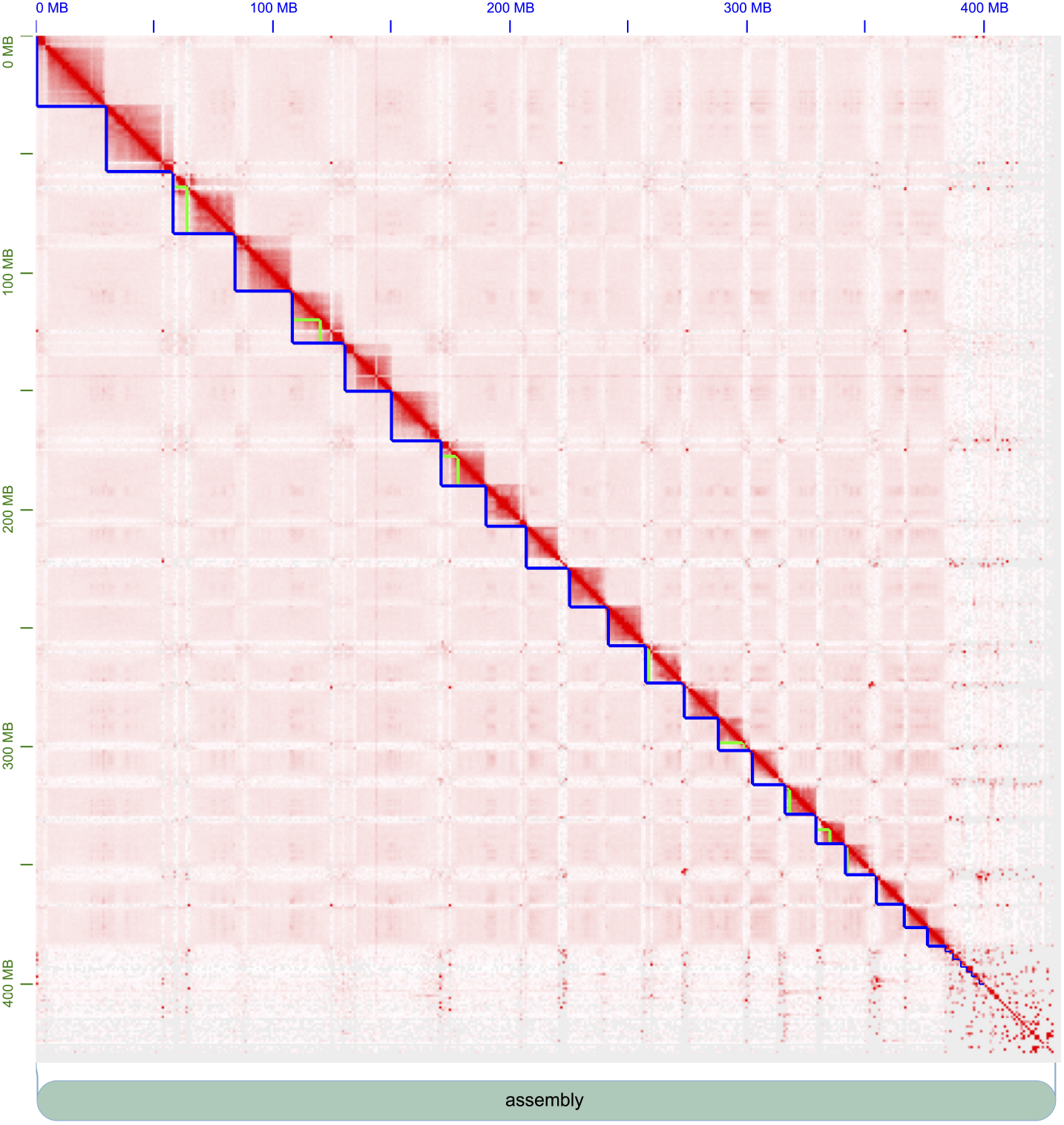
Visualization of contact maps from Hi-C reads for *Syngnathus scovelli*(v2). The first 22 primary assembly features (blue lines) sum to about 380Mb in size, which is the estimated genome size for the species. Green lines reflect individual contigs from the hifiasm assembly that were organized into chromosome-level scaffolds based on Hi-C contact data.

### BUSCO and Merqury Results

BUSCO results suggest a high degree of completeness as it finds 95% of the orthologs in the actinopterygii dataset (94.7%[*S* : 93.9%, *D* : 0.8%], *F* : 1.5%, *M* : 3.8%, *n* : 3640) when run in genome mode (Figure 2) and the Merqury evaluation suggests that the genome is ∼86% complete with a QV value of 61.37 and an error rate of 7.3*e*^−05^% (see supplementary for more details; Supplementary Table S2 and S3).

Consistent with the BUSCO contiguity metrics, the genome is on par with *S. acus* for completeness, which is also around 95% complete. Missing genes make up the majority of the remaining 5% of genes. We identify genes likely to be truly missing from the *S. scovelli* genome and more broadly among members of Syngnathidae (including the seahorses, genus *Hippocampus* along with *Syngnathus*) by confirming their absence across BUSCO results from the present assembly, four additional members of the genus *Syngnathus*, and six additional *Hippocampus* publicly available assemblies (see supplementary for additional details; Supplementary Table 6). Of the missing BUSCO genes, 83 genes are shared among all the species of *Syngnathus*, and 38 genes are missing from both genera (see supplementary for additional details; Supplementary Table 6). Future work could profitably explore these missing genes, as some may be related to the interesting novel traits in syngnathid fishes.

### Annotation Results

After masking about 43% of the genome, the annotations resulted in the prediction of about 28,162 genes, of which 8,061 were non-coding genes (see supplementary details; Supplementary Table 4 and Table 5). The 28,162 genes produce about 59,938 transcripts, of which 47,846 are mRNA, and the rest is made up of other types of RNAs such as tRNA, lncRNA, and others. Out of 20,101 coding genes, 18,616 genes had a protein with an alignment covering 50% or more of the query against the UniProtKB curated protein set, and 9152 had an alignment covering 95% or more of the query.

## Reuse Potential

The new version of the *S*.*scovelli* genome opens the doors for more accurate results by enhancing comparative genome data analysis and facilitating the creation of robust tools for molecular genetic studies. We generated the original version of the genome to focus on genetic mechanisms underlying the unique body plan among pipefishes and seahorses. This genome version takes us one step closer to uncovering these evolutionary mysteries and aids in answering other unknown features, such as the effects of sexual selection and mate choice systems on genome evolution.

## Data Availability

The genome is available on NCBI with the assembly accession number GCA_024217435.2. The genome is annotated via the NCBI eukaryotic genome annotation pipeline, and the annotation report release (100) is available here. Several smaller contigs and contaminant microbes were removed in the annotation pipeline yielding a more robust genome assembly. The sequence identifier for the chromosome-level scaffolds is available in the supplementary materials (see supplementary details; Supplementary Table 7). The NCBI Bioproject accession number is PRJNA851781, the raw Hi-Fi sequence accession is SRR19820733, the Hi-C sequence accession is SRR22219025, and for RNA-Seq sequence files from various tissues are SRR20438584 - SRR20438604.

## Declarations

### List of abbreviations

Mb: Mega basepair
Gb: Giga basepair
RNA: Ribo Nucleic Acid
DNA: Deoxyribo Nucleic Acid
BUSCO: Benchmarking Universal Single-Copy Orthologs
QUAST: Quality Assessment Tool
CCS: Circular Consensus Sequence
Hi-Fi: High-Fidelity
SMRT: Single Molecule Real Time
NCBI: National Center for Biotechnology Information

## Ethical Approval (optional)

Not applicable

## Consent for publication

Not applicable

## Author’s Contributions

The corresponding authors are Balan Ramesh and Adam Jones. Author contributions, described using the CASRAI CRedIT typology (http://casrai.org/credit), are as follows:

Conceptualization: BR, CMS, SB, WAC, AGJ; Methodology: BR, CMS, SB, BDJ, EB; Software: BR, CMS, HH, MC; Validation: BR, CMS; Formal Analysis: BR, CMS; Investigation: BR, CMS; Resources: MC, BDJ, EB, MM; Data Curation: BR, CMS, MC; Writing – Original Draft Preparation: BR, CMS, AGJ; Writing – Review & Editing: BR, CMS, AGJ; Visualization: BR, CMS; Supervision: WAC, AGJ; Project Administration: CMS, SB, WAC, AGJ; Funding Acquisition: WAC, AGJ;

## Acknowledgements

We are truly grateful for the dedicated efforts of Emily Rose and her students at Valdosta State University, who collected the *S*.*scovelli* samples crucial for this work (Florida Fish and Wildlife Conservation Commission Permit: SAL-17-0182-E, SAL-18-0182-E). We also thank Jeff Bishop and Tina Arredondo from the University of Oregon (UO) GC3F for library preparation and sequencing assistance. We want to acknowledge the staff at Phase Genomics for their helpful Hi-C technical support. We are grateful to Mike Coleman and Mark Allen for assisting with Talapas Supercomputer Cluster at UO and Benji Oswald with Research Computing and Data Services at UI. We greatly appreciate the support of the NCBI staff for the Eukaryotic Annotation Pipeline. We thank Jacelyn Shu for her pipefish illustrations. We are grateful for the helpful comments and review by Sven Winter and Yue Song on the manuscript.

## Supplementary Tables and Figures

**Figure S1.**
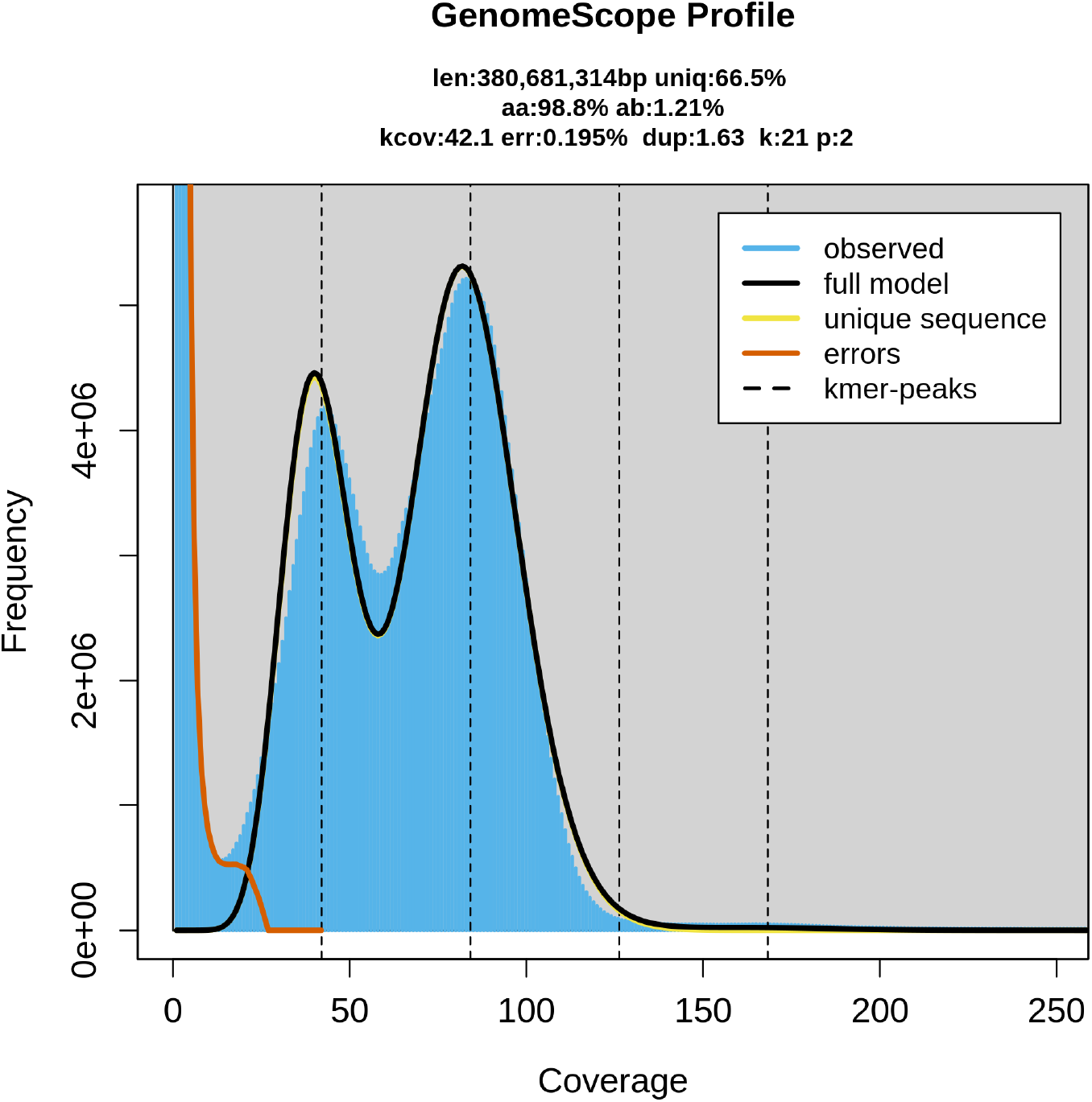
Genome Profile of *Syngnathus scovelli* based on Hi-Fi sequences using meryl and genomescope2 with k=21. The estimated genome size is around 380 Mb.

**Table S1.** Contiguity metrics from QUAST for the first pass and the scaffolded assembly of *S*.*scovelli*_v2 is summarized.

| Metrics | Haplotype1 | Haplotype2 | Primary Consensus Assembly | Scaffolded Assembly |
| --- | --- | --- | --- | --- |
| Number of contigs | 901 | 544 | 585 | 526 |
| Largest contig | 21,671,036 | 23,661,123 | 30,098,933 | 30,098,933 |
| Total length | 427,545,154 | 428,155,884 | 431,749,582 | 431,750,762 |
| GC (%) | 44.99 | 44.78 | 45.00 | 45.00 |
| N50 | 10,825,652 | 10,535,849 | 15,551,623 | 17,337,441 |
| N75 | 4,999,310 | 4,477,557 | 11,049,644 | 13,347,818 |
| L50 | 15 | 15 | 11 | 10 |
| L75 | 29 | 30 | 19 | 17 |
| Number of N's per 100 kbp | 0.00 | 0.00 | 0.00 | 0.27 |

**Table S2.** k-mer based assembly evaluation for completeness using Merqury

| Assembly | k-mer set used | solid k-mers in the assembly | Total solid k-mers in the read set | Completeness (%) |
| --- | --- | --- | --- | --- |
| *S.scovelli_v2* | all | 272,969,166 | 318,487,563 | 85.708 |

**Table S3.** k-mer based quality evaluation using Merqury

| Assembly | k-mers uniquely found only in the assembly | k-mers found in both assembly and the read set | QV | Error rate |
| --- | --- | --- | --- | --- |
| *S.scovelli_v2* | 6,614 | 431,737,882 | 61.3697 | 7.29504e-07 |

**Table S4.**
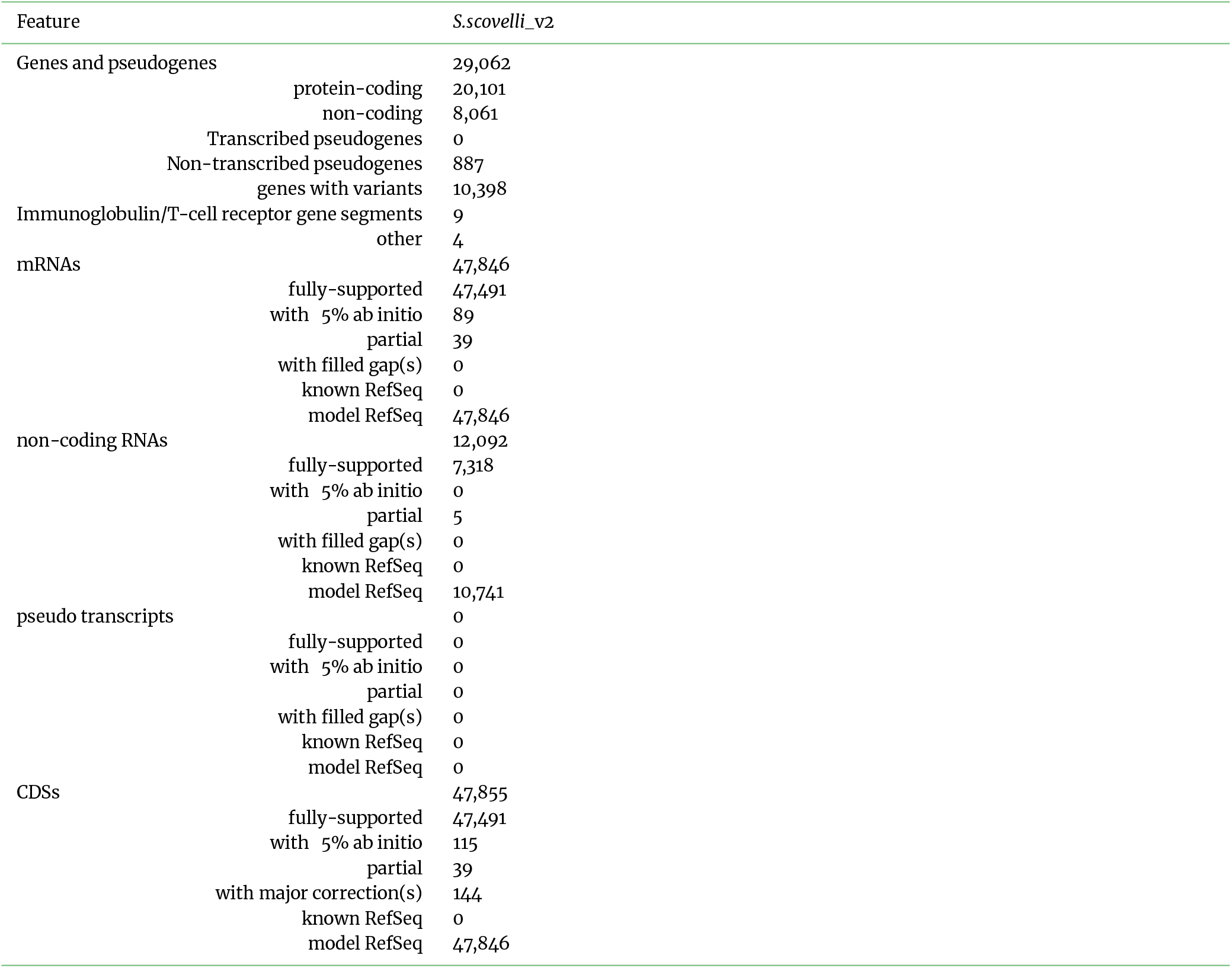
Gene and Feature Statistics from NCBI Eukaryotic Pipeline

**Table S5.** Detailed Feature Lengths from NCBI Eukaryotic Pipeline

| Feature | Count | Mean length (bp) | Median length (bp) | Min length (bp) | Max length (bp) |
| --- | --- | --- | --- | --- | --- |
| Genes | 28,166 | 11,149 | 4,361 | 56 | 677,970 |
| All transcripts | 59,938 | 3,654 | 2,773 | 56 | 106,526 |
| mRNA | 47,846 | 3,907 | 3,042 | 204 | 98,797 |
| miscRNA | 2,018 | 3,844 | 2,824 | 138 | 22,974 |
| tRNA | 1,351 | 74 | 73 | 71 | 87 |
| lncRNA | 5,304 | 3,880 | 1,632 | 112 | 106,526 |
| snoRNA | 117 | 123 | 126 | 62 | 319 |
| snRNA | 378 | 142 | 141 | 56 | 196 |
| rRNA | 2,920 | 1,228 | 154 | 118 | 4,380 |
| Single-exon | 514 | 2,381 | 1,944 | 358 | 21,617 |
| coding | 514 | 2,381 | 1,944 | 358 | 21,617 |
| CDSs | 47,846 | 2,373 | 1,617 | 96 | 97,746 |
| Exons | 277,161 | 325 | 142 | 2 | 38,823 |
| coding | 260,368 | 299 | 140 | 2 | 38,823 |
| non-coding | 27,774 | 515 | 152 | 9 | 36,521 |
| Introns | 247,597 | 1,355 | 160 | 30 | 611,280 |
| coding | 235,861 | 1,207 | 152 | 30 | 611,280 |
| non-coding | 22,579 | 2,911 | 304 | 30 | 498,241 |

**Table S6:** List of BUSCO IDS missing from odb10 in Syngnathidae

| S.No | Missing in *Syngnathus* genus | Missing in *Hippocampus* genus | Common among the two genera (*Syngnathus* and *Hippocampus*) |
| --- | --- | --- | --- |
| 1 | 57763at7898 | 67882at7898 | 55833at789 |
| 2 | 113518at7898 | 131304at7898 | 58251at789 |
| 3 | 70475at7898 | 65212at7898 | 129524at789 |
| 4 | 11981at7898 | 39050at7898 | 30261at789 |
| 5 | 35030at7898 | 55416at7898 | 69518at789 |
| 6 | 55416at7898 | 38468at7898 | 42298at789 |
| 7 | 80648at7898 | 37804at7898 | 92290at789 |
| 8 | 55833at7898 | 58251at7898 | 78658at789 |
| 9 | 15950at7898 | 43179at7898 | 50407at789 |
| 10 | 92290at7898 | 130413at7898 | 65212at789 |
| 11 | 75447at7898 | 88107at7898 | 106707at789 |
| 12 | 131401at7898 | 106707at7898 | 88107at789 |
| 13 | 73791at7898 | 55833at7898 | 59480at789 |
| 14 | 59450at7898 | 129524at7898 | 43179at789 |
| 15 | 65007at7898 | 107354at7898 | 37804at789 |
| 16 | 134401at7898 | 88998at7898 | 53002at789 |
| 17 | 85487at7898 | 75955at7898 | 103414at789 |
| 18 | 43533at7898 | 87291at7898 | 134401at789 |
| 19 | 1155at7898 | 107825at7898 | 88998at789 |
| 20 | 75955at7898 | 1155at7898 | 131304at789 |
| 21 | 30911at7898 | 102349at7898 | 104401at789 |
| 22 | 42232at7898 | 59450at7898 | 10242at789 |
| 23 | 71271at7898 | 8271at7898 | 38468at789 |
| 24 | 27668at7898 | 121548at7898 | 67882at789 |
| 25 | 10143at7898 | 53002at7898 | 10143at789 |
| 26 | 52590at7898 | 134401at7898 | 102349at789 |
| 27 | 86250at7898 | 69518at7898 | 21496at789 |
| 28 | 42298at7898 | 116at7898 | 107825at789 |
| 29 | 118274at7898 | 29564at7898 | 9176at789 |
| 30 | 58251at7898 | 42298at7898 | 59450at789 |
| 31 | 85116at7898 | 10143at7898 | 1155at789 |
| 32 | 43649at7898 | 92290at7898 | 75955at789 |
| 33 | 106801at7898 | 50407at7898 | 130413at789 |
| 34 | 82299at7898 | 103414at7898 | 57763at789 |
| 35 | 69518at7898 | 104401at7898 | 29564at789 |
| 36 | 103414at7898 | 9176at7898 | 55416at789 |
| 37 | 129524at7898 | 30261at7898 | 92234at789 |
| 38 | 110670at7898 | 59480at7898 | 93458at789 |
| 39 | 59480at7898 | 57763at7898 |  |
| 40 | 83539at7898 | 93458at7898 |  |
| 41 | 130413at7898 | 102866at7898 |  |
| 42 | 43179at7898 | 78658at7898 |  |
| 43 | 47247at7898 | 10242at7898 |  |
| 44 | 102349at7898 | 92234at7898 |  |
| 45 | 38468at7898 | 21496at7898 |  |

Table S6: List of BUSCO IDS missing from odb10 in Syngnathidae (Continued)
| S.No | Missing in *Syngnathus* genus | Missing in *Hippocampus* genus | Common among the two genera (*Syngnathus* and *Hippocampus*) |
| --- | --- | --- | --- |
| 46 | 88998at7898 |  |  |
| 47 | 106707at7898 |  |  |
| 48 | 92234at7898 |  |  |
| 49 | 51064at7898 |  |  |
| 50 | 43538at7898 |  |  |
| 51 | 111533at7898 |  |  |
| 52 | 107825at7898 |  |  |
| 53 | 9176at7898 |  |  |
| 54 | 14734at7898 |  |  |
| 55 | 15828at7898 |  |  |
| 56 | 10242at7898 |  |  |
| 57 | 78658at7898 |  |  |
| 58 | 93458at7898 |  |  |
| 59 | 109207at7898 |  |  |
| 60 | 69656at7898 |  |  |
| 61 | 56979at7898 |  |  |
| 62 | 104401at7898 |  |  |
| 63 | 79338at7898 |  |  |
| 64 | 88107at7898 |  |  |
| 65 | 97002at7898 |  |  |
| 66 | 122116at7898 |  |  |
| 67 | 60389at7898 |  |  |
| 68 | 116288at7898 |  |  |
| 69 | 131304at7898 |  |  |
| 70 | 67882at7898 |  |  |
| 71 | 70206at7898 |  |  |
| 72 | 65212at7898 |  |  |
| 73 | 91225at7898 |  |  |
| 74 | 50407at7898 |  |  |
| 75 | 53002at7898 |  |  |
| 76 | 84014at7898 |  |  |
| 77 | 30261at7898 |  |  |
| 78 | 37804at7898 |  |  |
| 79 | 29564at7898 |  |  |
| 80 | 7429at7898 |  |  |
| 81 | 21496at7898 |  |  |
| 82 | 44544at7898 |  |  |
| 83 | 4528at7898 |  |  |

The *Syngnathus* genus in this table includes *S. acus*, *S. rostellatus*, *S. typhle*, and *S. floridae* and the *Hippocampus* genus includes *H.comes*, *H.abdominalis*, *Herectus*, *H.kuda*, *H.whitei*, and *H.zosteræ*

**Table S7.** Sequence Identifiers for Chromosome Level Scaffolds

| Sequence Identifier |
| --- |
| NW_026061189.1 |
| NW_026061295.1 |
| NW_026061314.1 |
| NW_026061274.1 |
| NW_026061284.1 |
| NW_026061305.1 |
| NW_026061253.1 |
| NW_026061242.1 |
| NW_026061496.1 |
| NW_026061652.1 |
| NW_026061232.1 |
| NW_026061263.1 |
| NW_026061190.1 |
| NW_026061674.1 |
| NW_026061221.1 |
| NW_026061211.1 |
| NW_026061391.1 |
| NW_026061296.1 |
| NW_026061200.1 |
| NW_026061641.1 |
| NW_026061603.1 |
| NW_026061663.1 |

